# Removal of alleles by genome editing – RAGE against the deleterious load

**DOI:** 10.1101/335497

**Authors:** Martin Johnsson, R Chris Gaynor, Janez Jenko, Gregor Gorjanc, Dirk-Jan de Koning, John M Hickey

## Abstract

**Background:** In this paper, we simulate deleterious load in an animal breeding program, and compare the efficiency of genome editing and selection for decreasing load. Deleterious variants can be identified by bioinformatics screening methods that use sequence conservation and biological prior information about protein function. Once deleterious variants have been identified, how can they be used in breeding?

**Results:** We simulated a closed animal breeding population subject to both natural selection against deleterious load and artificial selection for a quantitative trait representing the breeding goal. Deleterious load was polygenic and due to either codominant or recessive variants. We compared strategies for removal of deleterious alleles by genome editing (RAGE) to selection against carriers. Each strategy varied in how animals and variants were prioritized for editing or selection.

**Conclusions:** Genome editing of deleterious alleles reduces deleterious load, but requires simultaneous editing of multiple deleterious variants in the same sire to be effective when deleterious variants are recessive. In the short term, selection against carriers is a possible alternative to genome editing when variants are recessive. The dominance of deleterious variants affects both the efficiency of genome editing and selection against carriers, and which variant prioritization strategy is the most efficient. Our results suggest that in the future, there is the potential to use RAGE against deleterious load in animal breeding.

## Background

Deleterious load is an unavoidable fact of genetics with a sizeable impact on the fitness of populations [1]. Most individuals have *de novo* deleterious mutations due to errors in DNA replication [2–4] and inherit many more from their ancestors. In this paper, we simulate deleterious load in an animal breeding program, and compare the efficiency of genome editing and selection for decreasing it.

Deleterious variants can have large or small effects. Recessive lethal variants are the most obvious symptoms of large effect deleterious mutations [5–12]. However, estimated distributions of deleterious mutation effects from several species indicate that most deleterious load is due to many variants of individually small effects [13–16]. In classic equilibrium models of deleterious variation [1], the effect size does not matter to the total load; only the mutation rate matters. In practice, however, large effect variants are easier to identify and manage. This raises the question: What can be done about polygenic deleterious load?

Deleterious variants of large and small effect can be identified by bioinformatics screening methods that use sequence conservation and biological prior information about protein function [17–21]. Such approaches have been applied to whole-genome sequence data to detect deleterious variants in crop plants [22–24], livestock [25–27], and humans [28,29]. With the decreasing cost to genome sequencing, and large initiatives to sequence livestock animals, we can anticipate sequence variant screening as a routine part of animal breeding.

Once deleterious variants have been discovered, there are two obvious ways to incorporate them into breeding: genome editing or selection. Genome editing is a suite of methods to modify the genomic DNA of an organism, allowing not just insertion and deletion but replacement of sequences with higher efficiency than homologous recombination (reviewed by [30]). Genome editing has shown theoretical promise for improving breeding progress by promoting favorable alleles [31], and for managing recessive lethal variants [32]. Selection against carriers is the strategy of choice for removing monogenic recessive deleterious variants from animal breeding populations [5,33]. Analogously, one could select against deleterious load by avoiding selection candidates with high deleterious load.

The aim of this paper was to compare the efficiency of genome editing and selection against carriers for decreasing deleterious load in an animal breeding program. We simulated polygenic deleterious load subject to natural selection in a simulation of a closed animal breeding population artificially selected for a quantitative performance trait representing the breeding goal. We compared removal of alleles by genome editing (RAGE) to selection against carriers using genotypes at deleterious variants. We compared strategies for prioritizing variants for editing and individuals for selection based on deleterious allele and genotype frequencies. Our results showed that RAGE reduces deleterious load, but requires simultaneous editing of multiple deleterious variants in the same sire to be effective when deleterious variants are recessive. In the short term, selection against carriers is a possible alternative to genome editing when variants are recessive, but in the future, RAGE against the deleterious load has great potential in animal breeding.

## Methods

We used simulations to compare genome editing and selection against carriers using genotypes at deleterious variants. We simulated artificial selection for a quantitative trait representing the breeding goal, and natural selection for a fitness trait representing reduced probability of survival due to deleterious variants. The fitness trait was polygenic with multiplicative fitness effects and an effect size distribution that was inspired by estimates of the distribution of deleterious effects in human populations [13].

In summary, the simulations consisted of 50 replicates of:

1. coalescent process simulation to create ancestral haplotypes;
2. setting up a quantitative trait and a fitness trait;
3. 15 generations of natural selection against deleterious variants, the first 5 using 1000 random matings per generation and the following 10 using 500 random matings per generation;
4. 20 generations of historical breeding with natural selection and simultaneous selection on true breeding value for the breeding goal trait; and finally
5. 10 generations of future breeding, where we evaluated scenarios with genome editing or selection against carriers.

### Simulation of whole genome sequence data and historical evolution

We used the Markovian Coalescent Simulator [34] to generate ancestral haplotypes. We modelled a genome consisting of ten chromosomes each of one Morgan in length with 6.75 × 10^8^ basepairs. The chromosomes were simulated using a per site mutation rate of 1.6 × 10^−8^, and an effective population size that changed over time to final size of 100. The effective population size was set to be 10^6^ at 190,000 generations ago, 100,000 at 100,000 generations ago, and 100 at current time, with linear decreases (on the 4 × N_e_ × time scale) in between.

### Simulation of quantitative and fitness traits

To capture artificial selection for the breeding goal and natural selection against deleterious variants simultaneously, we modelled a quantitative breeding goal trait and a fitness trait.

The breeding goal trait was a polygenic quantitative trait with additive effects. We randomly assigned 10 000 segregating sites (1000 per chromosome) as quantitative trait variants for the breeding goal trait with additive effects drawn from a normal distribution.

Fitness was a polygenic multiplicative trait that represented probability of survival prior to artificial selection. We randomly assigned 10 000 segregating sites as fitness variants (again 1000 per chromosome), choosing variants that with allele frequencies below 0.01 for codominant variants and 0.1 for recessive variants. The fitness variants were chosen independently of the quantitative trait variants. The deleterious effect size was expressed as a selection coefficient s against the mutant allele, going from 0 (no deleterious effect) to 1 (a lethal allele). The fitness of each genotype was *1* for the homozygous wildtype, *1 – h s* for the heterozygote, and *1 – s* for the mutant homozygote, where h is a dominance coefficient.

Dominance coefficients were either 0 for recessive variants or 0.5 codominant variants. We assumed multiplicative effects, so that the fitness of an individual was the product of the contribution of each fitness variant. The effect sizes were drawn from a mixture of three uniform distributions with one third of variants being small (0 < s < 10^−4^), one third intermediate (10^−4^ < s < 0.1), and one third large (0.1 < s < 1). These proportions were chosen based on the estimated distribution of deleterious effects in humans [13].

Deleterious mutations occurred randomly during burn-in and historical breeding with a per locus mutation rate of 10^−4^, to give a deleterious mutation rate of 1 per individual and genome. This is a conservative estimate for the deleterious mutation rate in mammals. No back-mutation was allowed, meaning that only wild type alleles could mutate. Quantitative trait variants for the breeding goal trait did not mutate, except during the initial coalescent simulation to create ancestral haplotypes.

### Pedigree structure and selection for the breeding goal trait

At each generation during the historical and future breeding, we first applied natural selection for fitness, then artificial selection for the breeding goal trait on the remaining individuals. For natural selection, we drew a uniformly distributed random number between 0 and 1 for each individual. If the number was greater than the fitness value for that individual, the individual was removed from the population before selection. For artificial selection, we selected 25 sires and a variable number of dams based on true breeding value for the breeding goal trait. Mating between sires and dams was random. Each dam had 10 progeny. The number of dams selected was based on the number of dams required to give a population of 5000 individuals at the average level of deleterious load at the end of the burn-in phase.

### Deleterious variant discovery

To simulate deleterious variant discovery, we selected a random fraction of the deleterious variants that segregated at the end of historical breeding and assumed them to be discovered. We used a discovery rate of 0.75 for the main scenarios, but also tested discovery rates of 0.5 and 1. To simulate imperfect detection of deleterious variants, we chose neutral segregating variants as false positives at random. We added false positives so that the total number of variants detected was equal to the number of segregating deleterious variants, and if discovery rate was d, a fraction 1-d were false positives. These discovered variants were allowed to be edited or used for selection against carriers subsequently.

### Scenarios

#### Removal of deleterious alleles by genome editing (RAGE)

During future breeding, we removed alleles by genome editing at discovered deleterious variants in all the selected sires. For variants where a sire was not already homozygous wildtype, we edited the genotype to homozygous wildtype, until a set number of variants had been edited. We assumed that editing was accurate in that it always produced wild type homozygotes, and had no deleterious off target effects. We edited 1, 5, or 20 variants per sire. We only edited variants that were discovered and segregating in the population.

We used five different strategies for prioritizing variants for editing. These strategies were based on information that would be available from genotyping the sires at discovered deleterious variants, namely the deleterious allele and genotype frequencies. We assumed that the deleterious effect size was unknown. The strategies were:

- By high frequency, removing variants in decreasing order of deleterious allele frequency.
- By low frequency, removing variants in increasing order of deleterious allele frequency.
- By lack of homozygotes, removing variants in decreasing order of the difference between observed and expected deleterious allele homozygotes.
- By intermediate frequency, removing variants in decreasing order of deleterious allele frequency after applying a threshold to exclude variants with allele frequency higher than 0.25.
- Random, in random order, using the same random order for all sires.

For comparison, we also ran a baseline scenario without genome editing, starting from the same initial populations after historical breeding.

#### Selection against carriers

During future breeding, we performed selection against carriers in sires by identifying carriers of high deleterious load and removing them before selection. We avoided the 100, 250, or 500 individuals with the highest load when selecting sires.

We used three different strategies for selecting carriers. These strategies were based on information that would be available from genotyping the sires at discovered deleterious variants, namely the deleterious allele frequencies, genotype frequencies, and individual numbers of deleterious alleles. The strategies were:

- Total load, avoiding individuals carrying the greatest number of deleterious alleles, summing over the discovered variants.
- Heterozygous load, avoiding individuals carrying the greatest number of deleterious alleles in a heterozygote state.
- Homozygous load, avoiding individuals carrying the greatest number of deleterious alleles in a homozygous state.

For comparison, we also ran a baseline scenario without selection against carriers, starting from the same initial populations after historical breeding.

### Metrics and statistical analysis

We evaluated the simulated scenarios by the improvement in average fitness of the individuals in the population. By an individual’s fitness, we mean the genetic value for the fitness trait. We calculated the average change in fitness from the first to the tenth generation of future breeding, and compared it to the change in fitness in a baseline condition without genome editing and selection against carriers, reporting mean and standard error of the mean.

We evaluated the effect of total number of fitness variants in the genome and their dominance coefficients on the number of segregating variants, the frequencies of deleterious alleles, and the deleterious load. By number of segregating variants, we mean the number of fitness variants that remained variable in the population after burn-in and historical breeding. By deleterious load, we mean the number of deleterious alleles carried by an individual. When considered separately, heterozygous load means the number of deleterious alleles carried in a heterozygous state, and homozygous load means number of deleterious alleles carried in a homozygous state.

We performed simulations using AlphaSimR (Gaynor, in prep), modified to allow for fitness traits. AlphaSimR runs on the R statistical environment [35], and uses Rcpp and Armadillo [36–38]. We calculated summary statistics in the R statistical environment, and made graphs with ggplot2 [39]. The simulation scripts are available from https://bitbucket.org/hickeyjohnteam/rage/.

## Results

Our results show that both genome editing of deleterious alleles and selection against carriers can reduce deleterious load in some cases, but is inefficient in others. The efficiency of genome editing and selection against carriers, and which variant prioritization strategy is the most efficient, depends on whether the deleterious variants are codominant or recessive.

### Deleterious allele frequencies and load in simulated populations

We simulated closed animal breeding populations under selection for a breeding goal trait, simultaneously affected by deleterious load consisting of either codominant or recessive variants. After historical breeding, the simulated populations had on average 4411 segregating deleterious variants in the codominant case, and 3456 in the recessive case. Each individual carried a load of 53 deleterious alleles on average in the codominant case and 95 deleterious alleles in the recessive case. Figure 1 shows violin plots of the deleterious load carried by individuals at both levels of dominance. We also varied the total number of fitness variants in the genome to assess the sensitivity to this assumption. The resulting average fitness was similar (Additional file 1).

**Figure 1:**
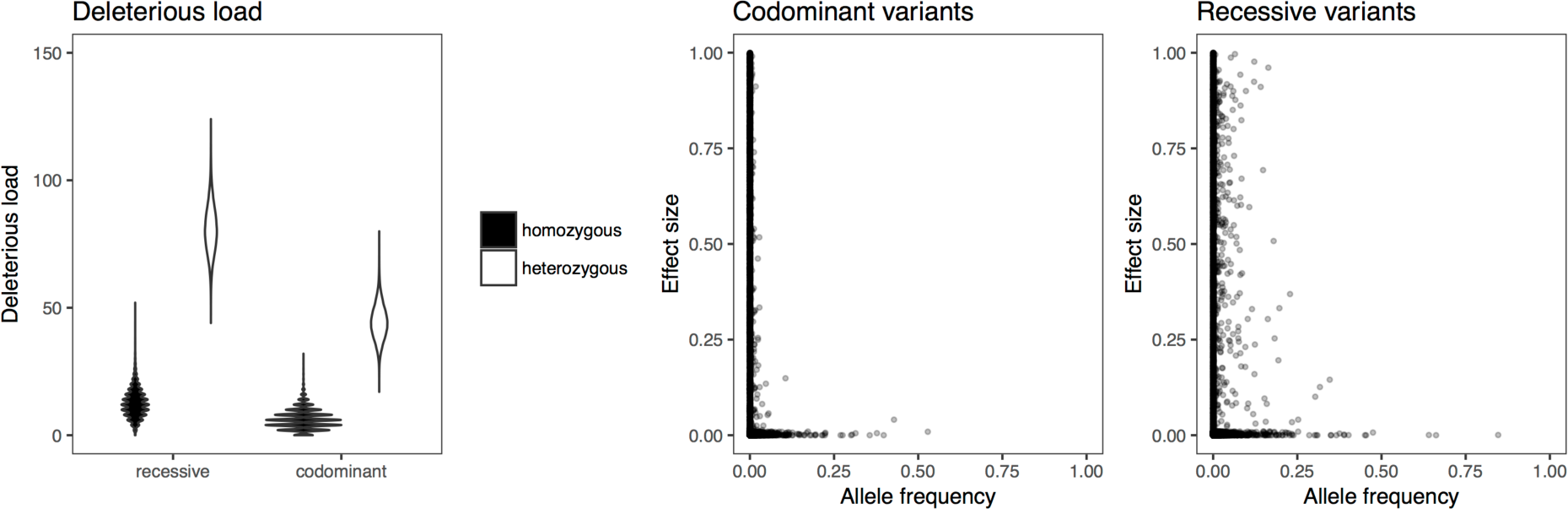
Deleterious allele frequencies and load in simulated populations when deleterious variants are either codominant or recessive. The violin plot shows individual deleterious load broken down into heterozygous and homozygous load with codominant or recessive variants. The scatterplots show the relationship between deleterious allele frequency and effect size.

Dominance also affected the distribution of frequencies of deleterious alleles. The scatterplots in Figure 1 show the relationship between effect size and frequency of deleterious alleles after historical breeding. When deleterious variants were codominant, most deleterious variants were rare. When deleterious variants were recessive, there was a larger number of deleterious alleles, even including large effect variants, at intermediate frequencies. These intermediate frequency deleterious alleles of large effects are candidates for removal by genome editing.

### Comparison of RAGE and selection against carriers

This difference in the distribution of frequencies and effects translates to differences in the efficiency of RAGE and selection against carriers. Figure 2 shows a comparison of genome editing using the best performing variant prioritization strategy, and selection against carriers using total deleterious load. When deleterious variants were codominant, the best performing strategy was prioritizing low frequency variants for removal by editing, and selection against carriers was inefficient. When deleterious variants were recessive, the best performing strategy was prioritizing intermediate frequency variants for removal by editing, and selection against carriers was comparable to genome editing of one variant per sire. However, multiplex editing of five variants per sire outperformed selection against carriers.

**Figure 2:**
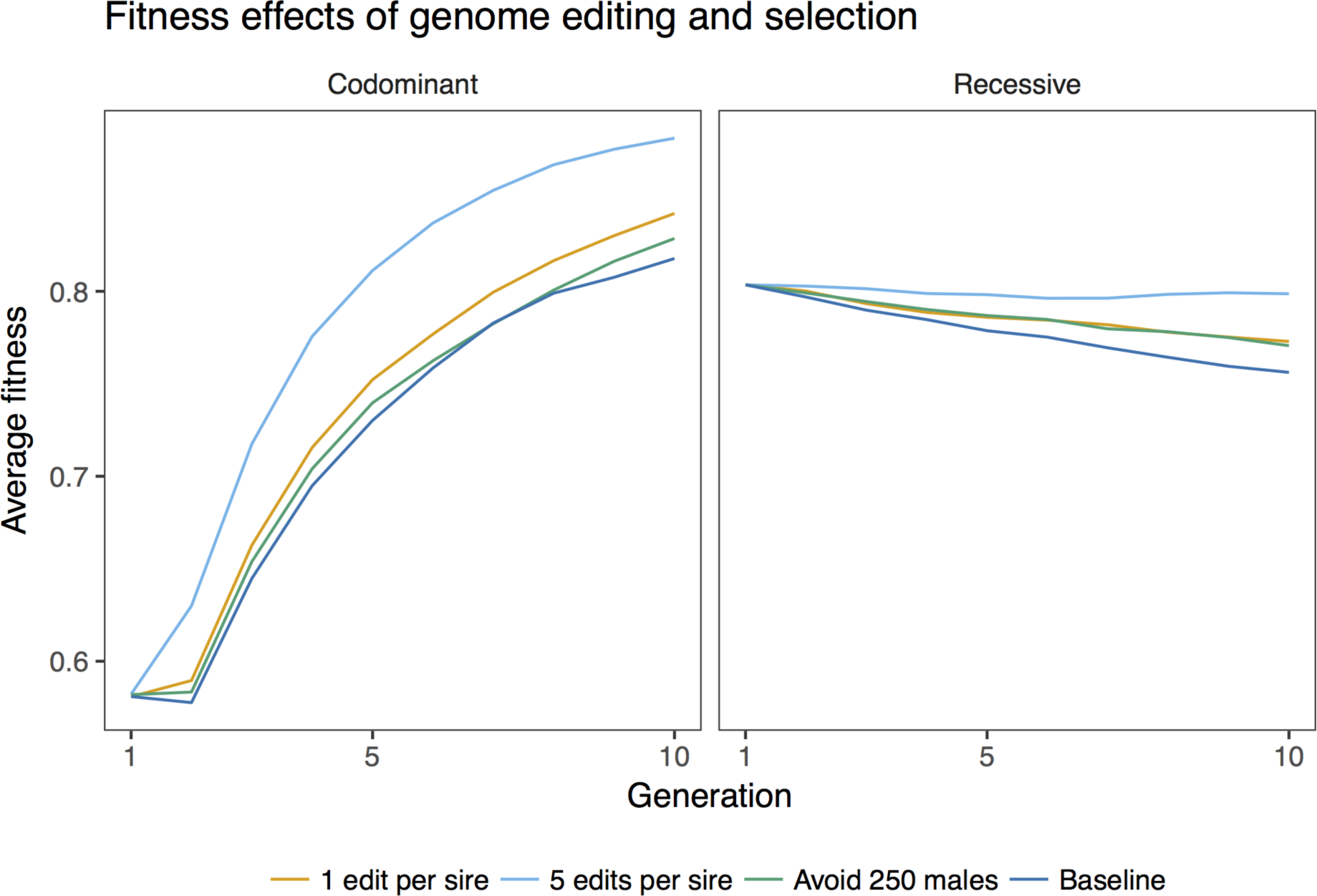
Comparison of the effect on average fitness of genome editing and selection against carriers, using the best performing editing strategies for each dominance level. The baseline condition is selection for the breeding goal trait with no effort to reduce deleterious load. The discovery rate was 0.75, meaning that 75% of the deleterious variants that segregated after historical breeding were discovered and could be edited. The lines show the average across 50 replicates.

We varied the number of variants edited per sire, the number of sires avoided, and the strategies for variant prioritization and selection in order to explore their effect on fitness improvement. Figure 3 shows the change in fitness under different scenarios after ten generations of future breeding, compared to the baseline case of breeding without genome editing or selection against carriers. In what follows, we present first the effects of different variant prioritization strategies on RAGE, then the effect of selection strategies on selection against carriers, and finally the impact of the ability to accurately detect deleterious variants.

**Figure 3:**
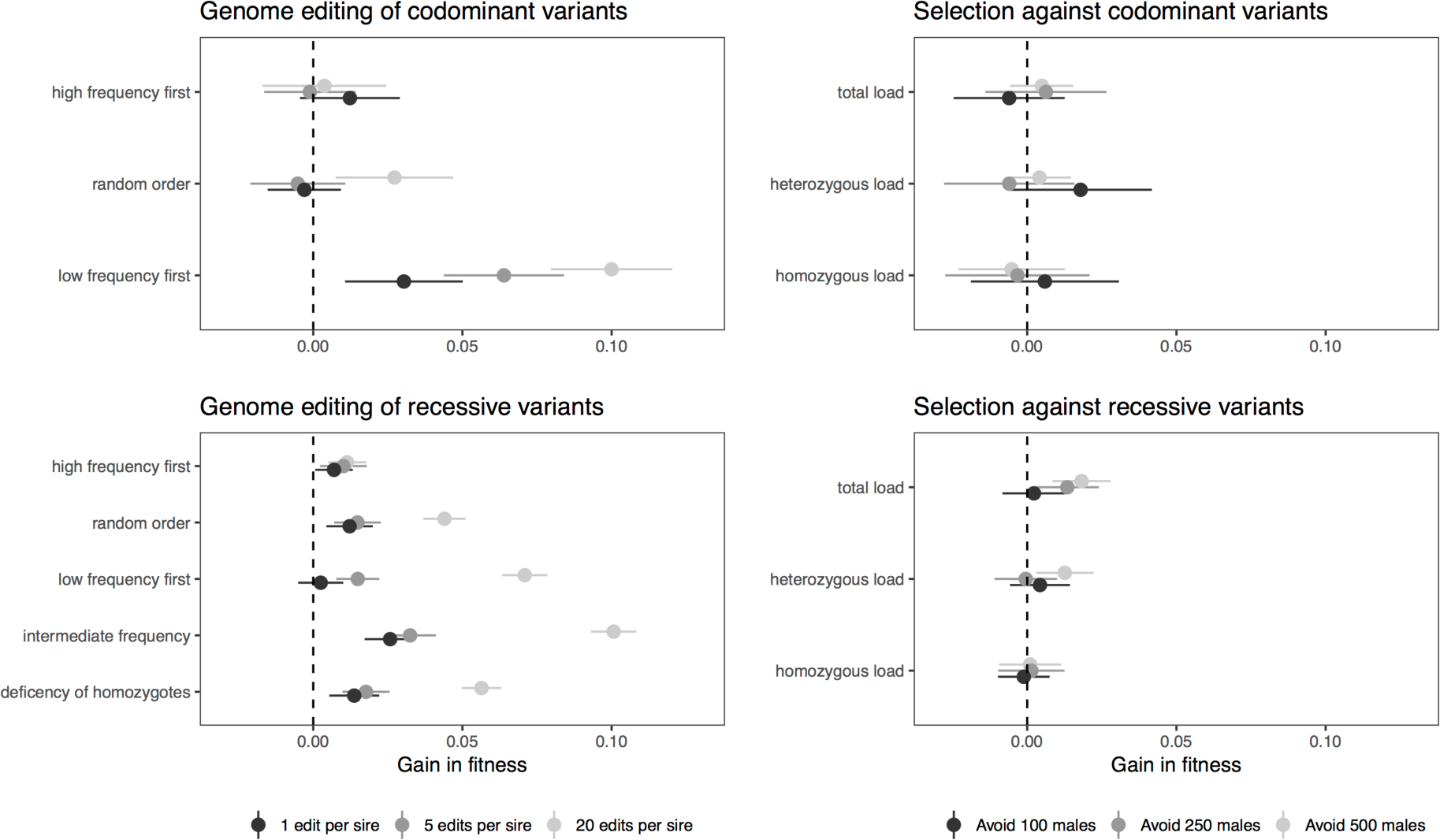
The effect on fitness of removal of deleterious alleles by genome editing and selection against carriers. The points show the mean change in fitness over ten generations compared to the baseline case of breeding without editing or selection, varying the number of edits per sire or the number of males avoided, and the strategy for variant prioritization or selection. The error bars are two standard errors of the mean.

Our model allowed population size to fluctuate with deleterious load, and scenarios using selection against carriers reduced the male population even further by excluding individuals with high load. This affected the genetic gain in the breeding goal trait by up to a 4% increase in gain in scenarios where deleterious load was alleviated, and up to a 5% loss of genetic gain with selection against carriers (Figure 4).

**Figure 4:**
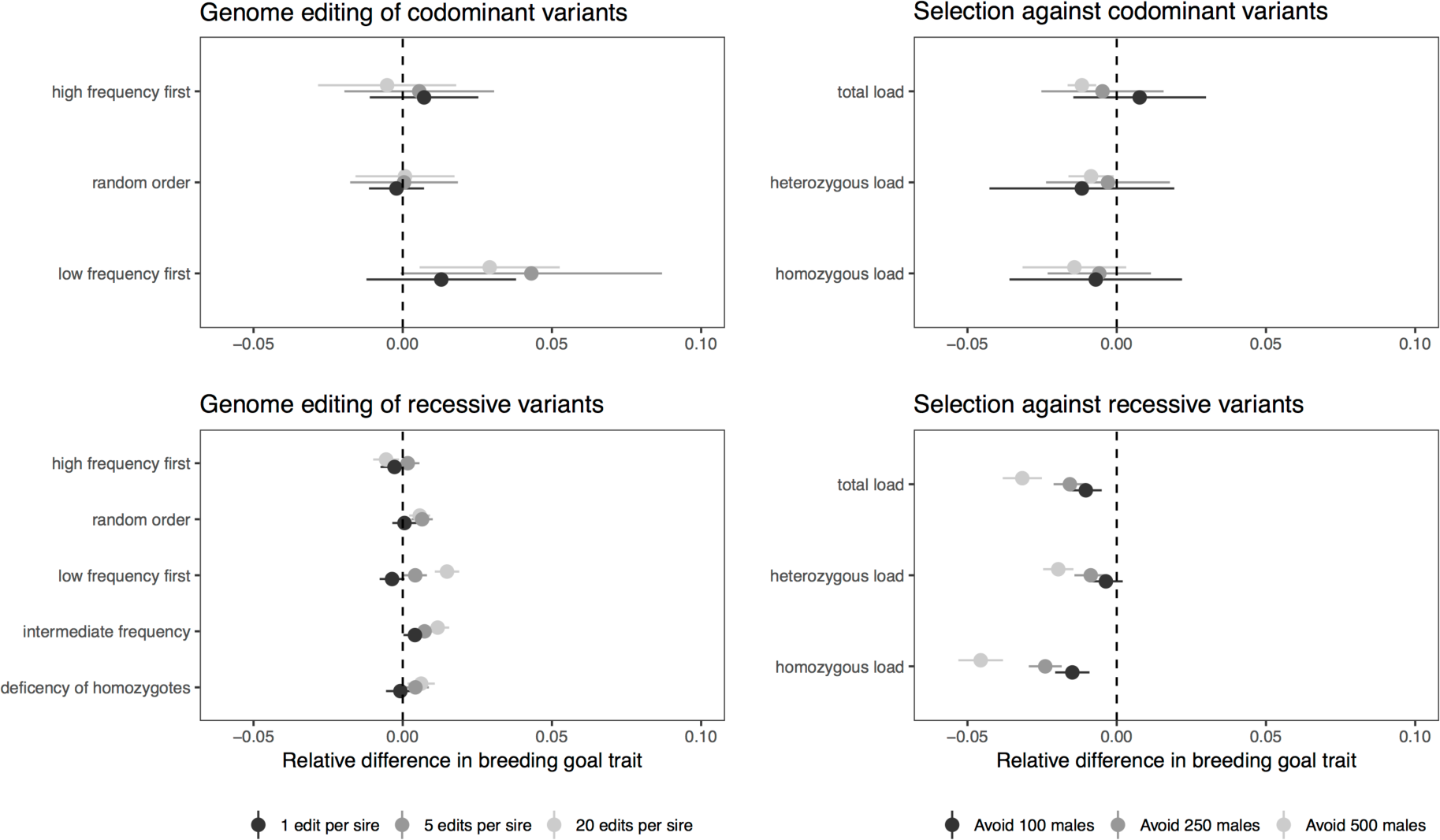
The effect on the breeding goal trait of removal of deleterious alleles by genome editing and selection against carriers. The points show the mean relative change in the breeding goal trait over ten generations compared to the baseline case of breeding without editing or selection, expressed as fraction of the increase without genome editing or selection against carriers. The error bars are two standard errors of the mean.

### Effect of variant prioritization strategy on RAGE

The efficiency of genome editing of deleterious variants was affected by the number of variants edited per sire, and the strategy for prioritizing variants for editing. Figure 5 shows trajectories of fitness over generations of genome editing, varying number of variants edited, and prioritizing low frequency, high frequency, or randomly chosen deleterious variants for editing. Figure 6 shows trajectories of fitness during future breeding using variant prioritization strategies devised for recessive variants: prioritizing variants with intermediate frequency by applying an allele frequency threshold of 0.25, and editing variants based on their deficit of homozygotes. In all these cases, the discovery rate was 0.75, meaning that 75% of segregating deleterious variants were discovered, and a 25% false positive rate.

**Figure 5:**
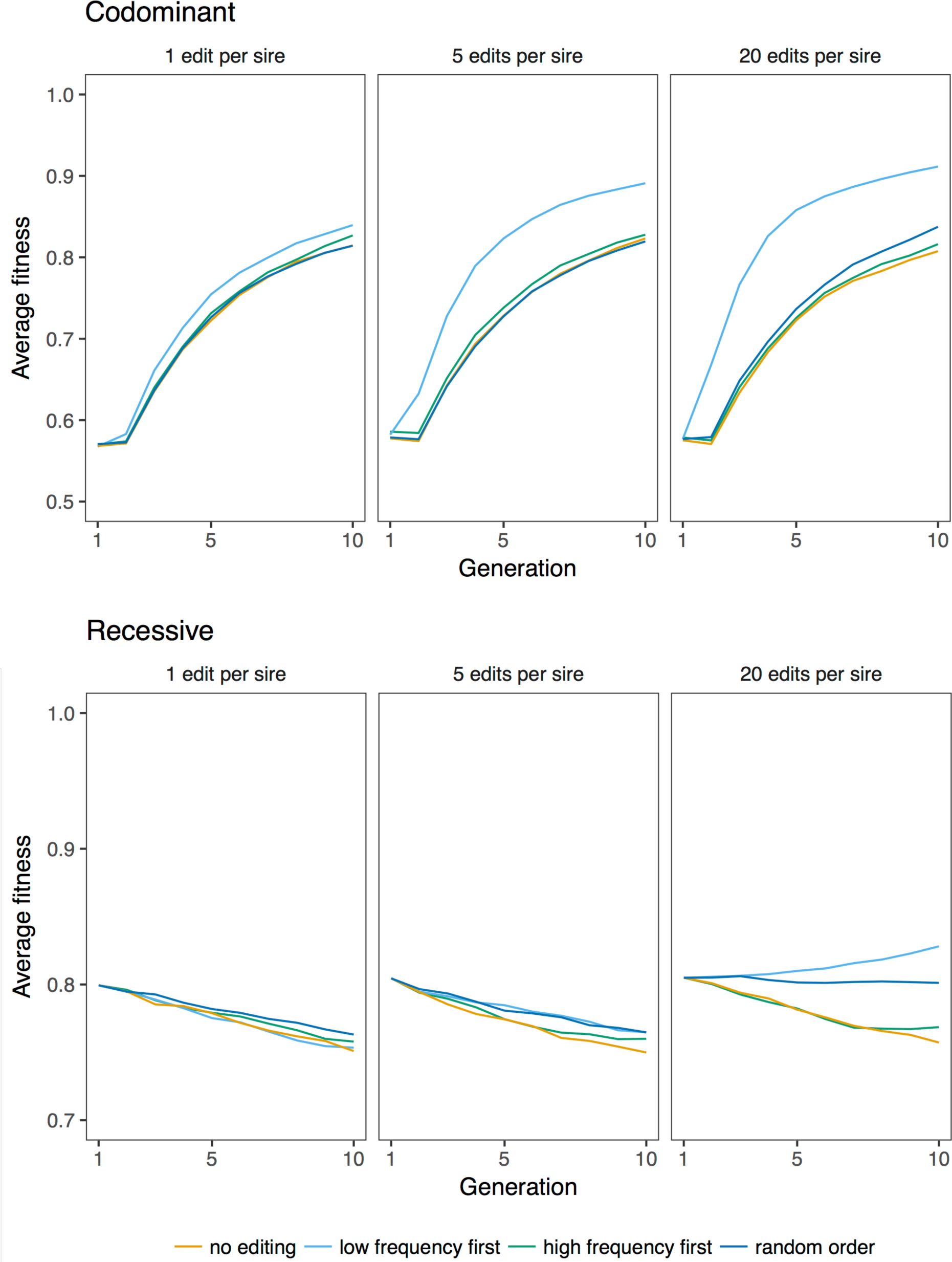
The effect of genome editing on fitness. Average fitness over ten generations of future breeding with different editing strategies, and editing of 1, 5, or 20 variants per sire. The discovery rate was 0.75. The lines show the average across 50 replicates.

**Figure 6.**
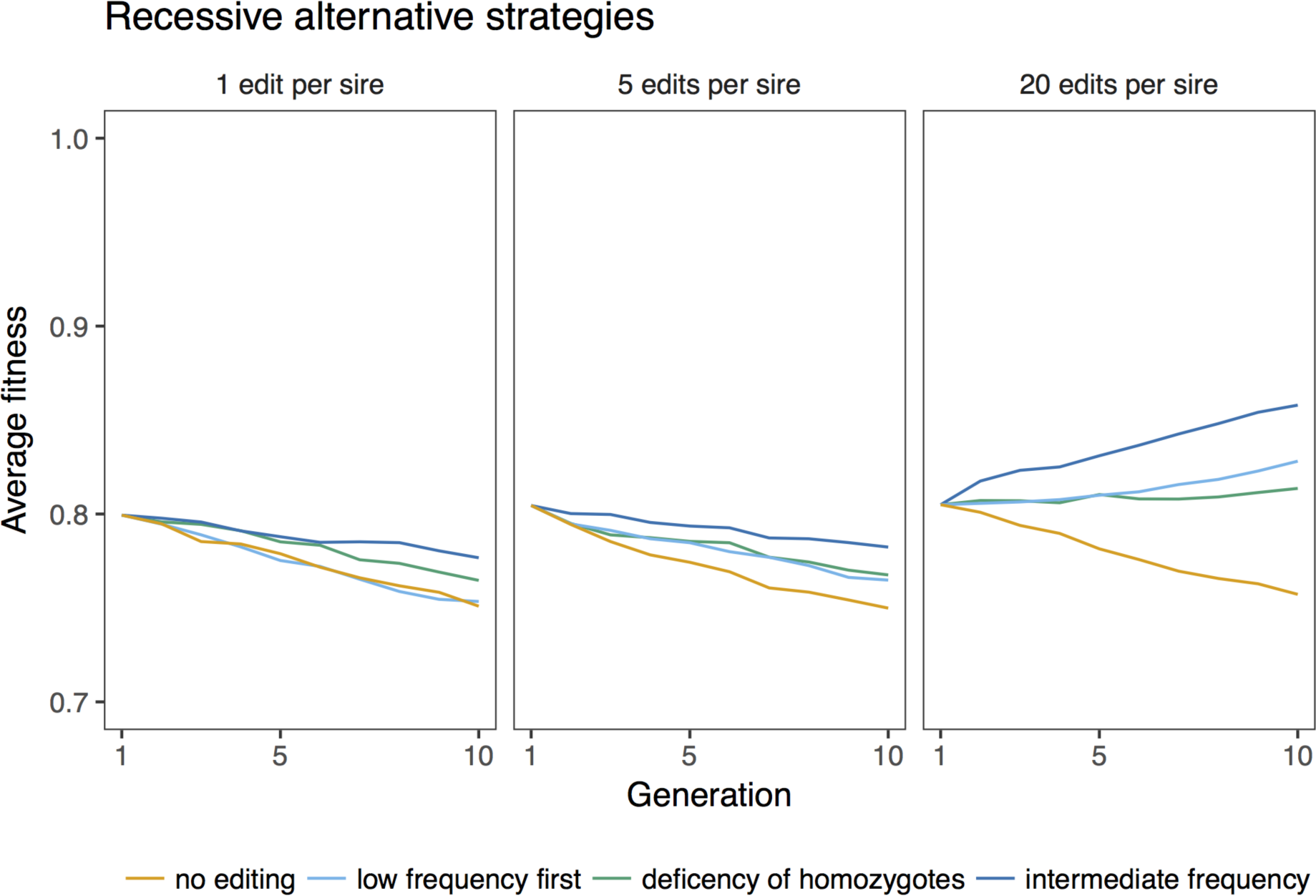
The effect of prioritizing variants with intermediate allele frequency, or in order of their deviation from expected homozygosity. Average fitness over ten generations of future breeding with 10 000 deleterious variants, editing 1, 5, or 20 variants per sire. The discovery rate was 0.75. The lines show the average across 50 replicates.

For both levels of dominance, fitness improved more by prioritizing low frequency variants for editing than by editing in random order, or prioritizing high frequency variants. When variants were recessive, fitness improved the most by prioritizing variants with intermediate allele frequency. Prioritizing variants with a deficit of homozygotes did not improve efficiency compared to using allele frequency.

The different variant prioritization strategies also differed in how many distinct variants were edited. Table 1 shows the average number of distinct variants edited during ten generations of future breeding with genome editing. Prioritizing low frequency variants for editing led to the largest number of distinct variants being edited, the random order and intermediate allele frequency strategies were intermediate, and prioritizing high frequency variants for editing led to the smallest number of distinct variants being edited.

**Table 1.**
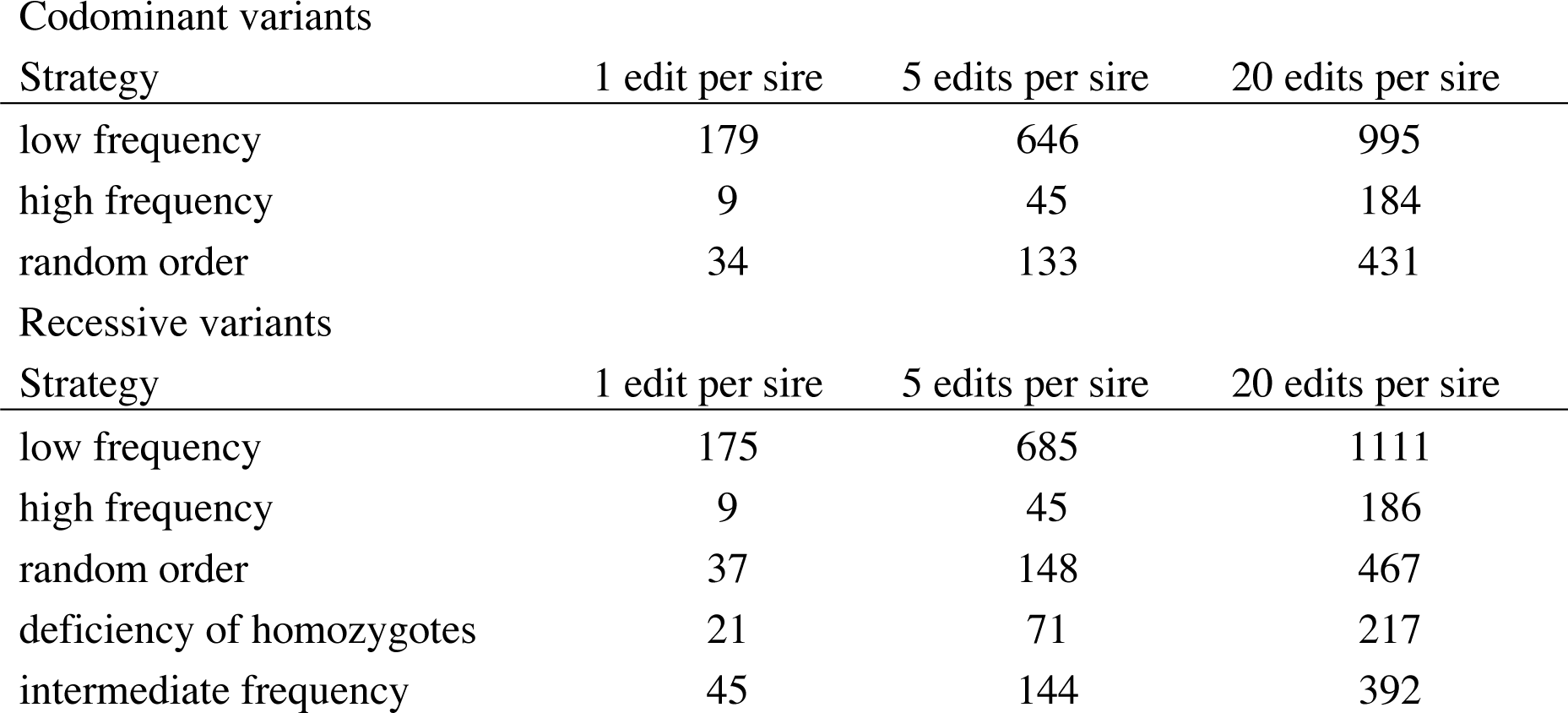
*Number of distinct variants edited over ten generations of future breeding with different strategies. The numbers are averages across 50 replicates.*

### Effect selection strategy on selection against carriers

The efficiency of selection strategies against carriers also varied with the number of males avoided, selection strategy, and dominance (Additional file 2). When deleterious variants were codominant, selection against carriers was inefficient regardless of strategy. When deleterious variants were recessive, selection on total load was the most efficient of the selection strategies. In no case was it better to select only on heterozygous or homozygous load.

### Effect of the ability to discover deleterious variants

The efficiency of genome editing and the relative performance of variant prioritization strategies was affected by the discovery rate. Figure 7 shows fitness trajectories when varying how many of the deleterious variants could be discovered and edited, showing an increase in fitness improvement with perfect discovery rate, but little difference between a discovery rate of 0.5 and 0.75, for all strategies. Thus, all strategies were susceptible to false positives, but it mattered less whether the false positives made up 25% or 50% of the detected variants.

**Figure 7:**
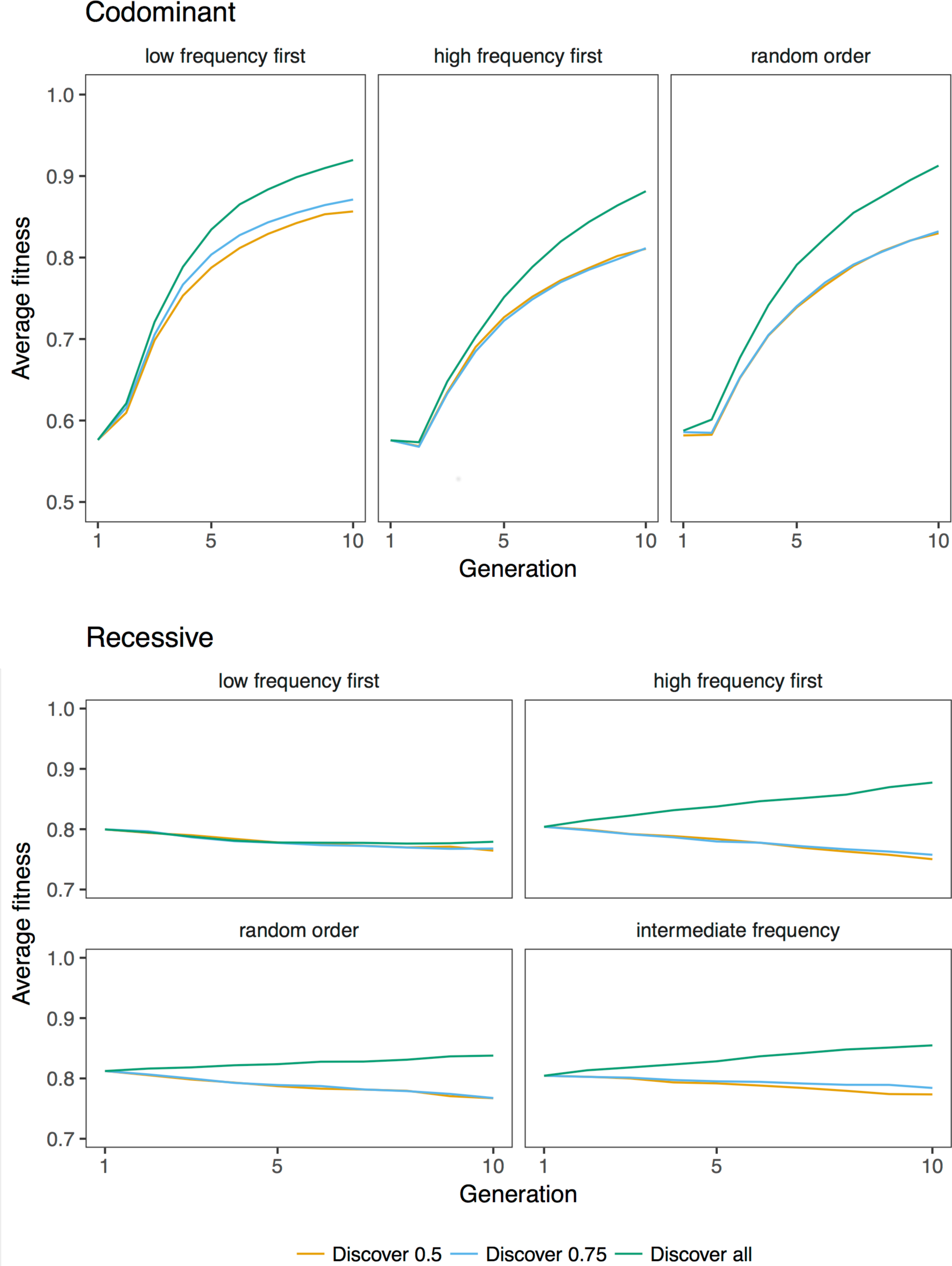
The effect of discovery rate on editing. Average fitness over ten generations of future breeding with 10 000 deleterious variants, 5 edited variants per sire, and discovery rates of 0.5, 0.75, and 1. The lines show the average across 50 replicates.

When variants were recessive, a high discovery rate changed the relative ranking of variant prioritization strategies: prioritizing high frequency variants for editing was more efficient than prioritizing intermediate frequency variants. Figure 8 shows fitness trajectories when all segregating deleterious variants were discovered with no false positives. Prioritizing intermediate frequency variants was the second-best strategy when one or five variants per sire were edited. When 20 variants per sire were edited, its efficiency plateaued, likely because this strategy involved excluding high frequency variants from consideration. Taken together, this means that the presence of false positives affects strategies that use allele frequency information for variant prioritization differently.

**Figure 8:**
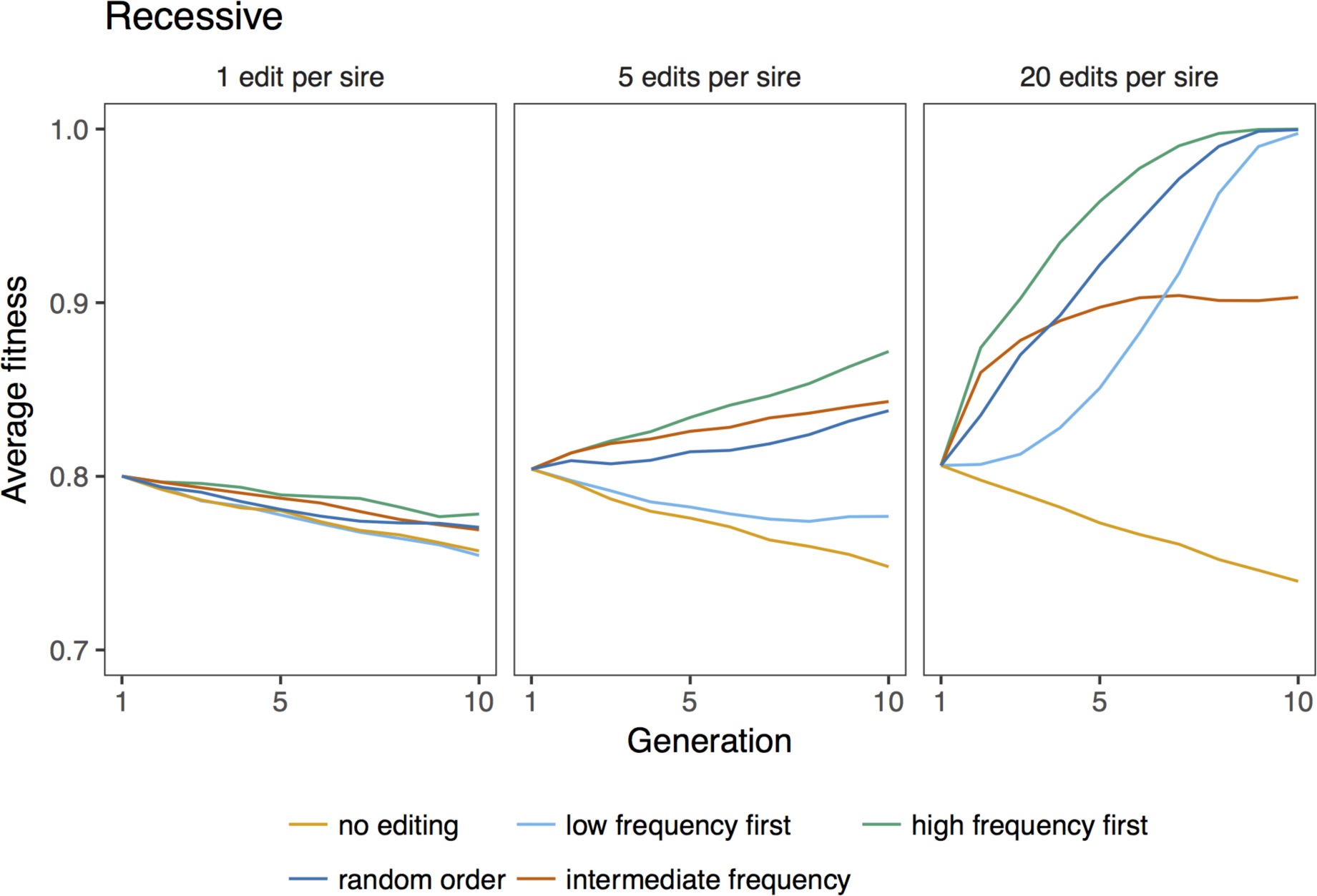
The relative efficiency of genome editing strategies against recessive deleterious variants when variant discovery is perfect (discovery rate is 1). The lines show the average across 50 replicates.

## Discussion

In this paper, we simulated deleterious load in an animal breeding program, and compared the efficiency of genome editing and selection for decreasing it. We found that both removal of alleles by genome editing and selection against carriers can reduce deleterious load in some scenarios. The dominance of deleterious variants affect the efficiency of genome editing and selection against carriers, and determines which variant prioritization and selection strategy is the most efficient.

In the light of these results, we will discuss (i) deleterious load in animal breeding populations, (ii) the efficiency of different variant prioritization and selection strategies, (iii) factors that will improve the efficiency of genome editing of deleterious variants, (iv) the assumptions underlying the simulations, (v) implications for applications of genome editing and selection against carriers in breeding, and (vi) open questions about the use of RAGE against deleterious load.

### Deleterious load in animal breeding populations

The deleterious loads in the simulated populations were comparable to observed loads of putative loss of function variants in mammals, but much smaller than the numbers of deleterious single nucleotide variants predicted with sequence bioinformatics-based methods. Humans are estimated to carry an average load of around 100 [40,41] or 150 [42] putative loss-of-function variants in protein-coding genes. The average load of loss-of-function variants observed in cattle is 65 [43]. The load of deleterious variants in our simulations (around 100 deleterious alleles per individual when variants were recessive) was comparable to these numbers. However, observed loads of deleterious nonsynonymous single nucleotide variants predicted by bioinformatics methods are much higher: around 300-800 for humans [44,45], and 656 in a pig population [46]. This suggests that our assumptions about the effect size distribution of deleterious variants or genomic deleterious mutation rate may be conservative, and the actual deleterious load may be higher.

### Efficiency of different variant prioritization and selection strategies

The dominance of deleterious variants affected the allele frequency distribution of deleterious alleles, and therefore determined which variant prioritization strategy and selection strategy was the most efficient.

When deleterious variants were codominant, large effect deleterious variants were rare. Because codominant deleterious alleles are expressed even when in a heterozygous state, they are exposed to purifying selection. Therefore, the best variant prioritization strategy was to prioritize low frequency variants for editing.

In contrast, when deleterious variants were recessive, there were substantial numbers of large effect variants at intermediate frequencies. This happens because of the inefficiency of natural selection against recessive variants. Because recessive deleterious variants are expressed only when in a homozygous state, they are more likely to do damage the more common they are. Despite this, in the presence of false positives, the best variant prioritization strategy was to prioritize intermediate frequency variants for editing. Because false discoveries will be neutral variants, they will on average have higher frequencies than genuine deleterious variants. Therefore, the high frequency variant prioritization strategy is especially susceptible to false positives. Prioritizing intermediate frequency variants for editing balances these effects, at least when the number of edits per sire are low. Another benefit of prioritizing intermediate frequency variants, or randomly selected variants, for editing is that this strategy requires fewer distinct variants to be edited, and therefore fewer proven editing constructs to be developed and tested, compared to prioritizing low frequency variants.

Our simulations showed no benefit from prioritizing recessive variants for editing based on deficit of homozygotes. The strategy was inspired by the method for discovering recessive lethal haplotypes of VanRaden et al [6], who compared the number of observed and expected homozygotes. This and related methods have been successfully used to detect recessive lethal haplotypes in livestock. Its failure as a variant prioritization method may be due to the fact that the simulated deleterious variants had variable effects, most of them individually of small effect, or the fact that many variants were rare and therefore would have low numbers of expected homozygotes. Furthermore, the frequency of lethal haplotypes may be exaggerated by extremely widespread use of a few sires in cattle populations [6].

In real populations, we should expect deleterious variants to be recessive, because they persist longer in the population. Therefore, we should prioritize editing deleterious variants with intermediate frequencies, or avoid carrier males based on their total deleterious load.

### Factors that will improve the efficiency of RAGE

According to our simulations, the most important factor for increasing the efficiency of removal of deleterious alleles by genome editing is the ability to edit multiple variants per individual. It is currently not feasible to produce germline edited livestock with multiple alleles edited, but genome editing technology is developing very rapidly. Multiplex genome editing has been performed in cells and model organisms [47–51].

The ability to accurately discover deleterious variants is also important. Methods to detect deleterious variants either predict variant consequences (e.g. stop codons and frame shifts) based on the genetic code [52,53], measure evolutionary conservation or constraint in multiple sequence alignments [17,19,20], or train statistical models to classify variants based on known deleterious variants and various predictors, possibly including variant consequences and evolutionary constraint, or functional genomic and protein structure data [18,21,28,54,55]. We expect the latter approach to become more accurate as machine learning methods improve and get access to ever bigger datasets of genetic variants and genomic data, and it becomes possible to train models on livestock rather than human data.

Our simulations assumed that there was no information on the effect size of the predicted deleterious variants. This is a conservative assumption, because in fact, it may be possible to stratify predicted deleterious variants by predicted impact. For example, protein-coding variants are likely to have greater effect sizes than non-coding variants [56,57], and loss-of-function variants are likely to have greater effects sizes than nonsynonymous single nucleotide variants.

### Assumptions underlying the simulations

Assumptions about the genetic architecture of deleterious load in these simulations include: the number of fitness variants in the genome, independent genetic architectures of the breeding goal trait and fitness, a genomic deleterious mutation rate of 1, and equal dominance coefficients for all variants. In real genomes, we expect that many more than 10 000 sites can give rise to deleterious mutations, but since the number of segregating variants was not much affected by the total number of fitness variants in the genome, this assumption appears to have little impact on results. We simulated fitness as independent of the selected performance trait. In real populations, we expect that fitness is to some extent already part of the breeding goal in the form of survival, fecundity, and health traits. This means that it is possibly to validate deleterious variants by phenotypic means, and including them in genomic selection models [24]. We assumed a genomic deleterious mutation rate of 1, but this is a conservative estimate, given that deleterious mutation rates for humans are often estimated to be higher (e.g. 1.6-3) [2–4]. We assumed equal dominance coefficients for all variants: either 0.5 (codominant) or 0 (recessive). In real populations, there could be a range of dominance coefficients, but recessive variants are expected to persist longer in the population.

### Implications for breeding

We found that genome editing of deleterious alleles reduced deleterious load, but that when variants were recessive, simultaneous editing of multiple deleterious variants in the same sire was needed for it to be competitive with selection against carriers. When accurate multiplex genome editing becomes available, RAGE has the potential to improve fitness to levels that are impossible by selection against carriers. This is a formidable undertaking, but a possible long term goal. The long-term benefits of genome editing to remove deleterious variants over selection against carriers include both the possibility of greater gains in fitness, and the ability to improve fitness without sacrificing selection intensity for the breeding goal trait.

In the short term, selection against carriers is a possible alternative to genome editing. It is ineffective against codominant variants, but when variants are recessive, it is more effective at alleviating deleterious load than editing one variant per sire, but it is less effective than multiplex editing. The cost of multiplex genome editing is unknown, but can assumed to be high. Therefore, it appears that selection against carriers will remain superior for some time. The downside of selection against carriers is that the number of sires available for selection is reduced, with associated risks of inbreeding and loss of genetic variation. Van Eenennaam and Kinghorn [58], and Cole [32,59] have extended mate selection schemes to also penalize the use of carrier animals. Possibly, such methods could be extended to penalize genome-wide deleterious load while maintaining diversity in other parts of the genome, and maximizing the response to selection for production traits.

To perform selection against carriers in practice, deleterious load would need to be included in a selection index and given an economic weight to balance it with the breeding goal, and make sure that selection against load does not unfavorably affect other traits. Unfavorable genetic correlations between deleterious load and traits could arise either from false positives, pleiotropy, or linkage disequilibrium. Deleterious variant prediction methods may mistakenly classify beneficial variants as deleterious because they change protein function. In fact, they may even have been deleterious in the wild, but beneficial in a modern farm environment, such as loss-of-function mutations in *myostatin* [60] that cause double muscling in beef cattle breeds. Deleterious variants may also have pleiotropic effects, or be found in linkage disequilibrium with selected variants. In all these cases, it may be possible to use marker estimates from genomic selection models to prune the deleterious variant set of variants associated with large beneficial effects on other traits before calculating deleterious load. Another option is to weight the variants by its associations to all the traits in a breeding goal and their hypothetical association with fitness to holistically balance their use in a breeding program.

### Open questions

Further research may find other ways to incorporate genomic information in variant prioritization for gene editing or in weighting of variants for selection against carriers. One genomic feature that may play into prioritization of deleterious variants for genome editing is recombination rate variation. In regions of low recombination, which in mammal genomes occur for example in centromeric regions and on the sex chromosomes, selection against deleterious variants is less efficient due to Hill—Robertson interference [61]. This phenomenon may both lead to accumulation of deleterious variants and reduced selection for beneficial variants that are located there. Therefore, it may also be beneficial to prioritize variants that experience low recombination rate for genome editing [62,63].

We have investigated removal of alleles by genome editing and selection against carriers to alleviate the load of deleterious variants that segregate within a population. Genome editing could also be used to remove deleterious alleles that are fixed in the population, and cannot be removed by selection. Fixed deleterious variants could be detected by sequencing studies that sample across populations and breeds that carry different sets of deleterious variants due to chance events such as mutation, genetic drift, and founder effects.

## Conclusions

Removal of alleles by genome editing reduces deleterious load, but requires simultaneous editing of multiple deleterious variants in the same sire to be effective. In the short term, carrier avoidance is a possible alternative to genome editing when variants are recessive. The dominance of deleterious variants affect the efficiency of genome editing selection and selection against carriers, and which variant prioritization strategy is the most efficient. Our results suggest that in the future, there is the potential to use RAGE against the deleterious load in animal breeding.

## Declarations

### Ethics approval and consent to participate

Not applicable.

### Consent for publication

Not applicable.

### Availability of data and materials

Data sharing is not applicable to this article as no datasets were generated or analyzed during the current study.

## Author’s contributions

JMH conceived the study. JMH and MJ designed the study. MJ performed the analysis. MJ and JMH wrote the manuscript. RCG, JJ, GG, and DdK helped interpret the result and refine the manuscript. All authors read and approved the final manuscript.

## Acknowledgements

This work has made use of the resources provided by the Edinburgh Compute and Data Facility (ECDF) (http://www.ecdf.ed.ac.uk).

